# Proximal Labeling of the Golgi Secretome Reveals Fat Body-Derived Humoral Factors in *Drosophila* Disc Regeneration

**DOI:** 10.1101/2024.09.02.609068

**Authors:** Yutaka Yoshida, Soshiro Kashio, Masayuki Miura

**Affiliations:** Department of Genetics, Graduate School of Pharmaceutical Sciences, The University of Tokyo, 7-3-1 Hongo, Bunkyo-ku, Tokyo 113-0033, Japan; Department of Integrative Bioanalytics, Institute of Development, Aging and Cancer (IDAC), Tohoku University, Sendai 980-8575, Japan; Laboratory for Cell Vigor Regulation, National Institute for Basic Biology, Nishigonaka 38, Okazaki, Aichi 444-8585, Japan

**Keywords:** Golgi, N-glycosylation, biotin labeling, proteomics, secreted protein

## Abstract

Humoral factors act as inter-tissue mediators and regulate various organismal physiologies. Although the importance of humoral factors in tissue repair has been recently recognized, our understanding of how humoral proteins regulate tissue repair remains limited. Glycosylation is an important modification of the conventional secretory pathway, and we have demonstrated that N-glycosylation in the Golgi apparatus of the *Drosophila* fat body (FB), the major secretory tissue equivalent to the mammalian liver and adipose tissue, remotely contributes to epithelial tissue repair. To identify humoral factors that contribute to repair via the Golgi apparatus, we constructed a Golgi-specific protein biotinylation system and performed hemolymph proteomics. By combining genetic analyses, we found that FB-derived innate immune regulators and iron mediators affect tissue repair. Altogether, our Golgi-specific labeling system has the potential to identify Golgi-mediated secreted factors that regulate inter-organ communication.

## Introduction

In multicellular organisms, the functions of each organ are highly specialized and coordinated. Various tools for analyzing inter-organ communication have recently emerged. In *in vivo* experimental systems, fluorescent-labeled exosomes are expressed in an organ-specific manner, and inter-organ communication was analyzed using zebrafish embryos (1). A method has also been developed to identify the trafficking of secreted proteins by protein biotin labeling in the endoplasmic reticulum (ER)(2).

Tissue regeneration is a homeostatic mechanism against tissue loss that is observed in specific organs and animals (3, 4). Non-autonomous tissue regulation of repair has also been examined. For example, the vagus-macrophage axis regulates liver regeneration in mice, and platelet-derived serotonin mediates liver regeneration (5, 6). *In vitro* screening showed that myristoylated alanine-rich C kinase substrate (MARCKS)-like protein, a factor secreted from the wound epidermis, is required for the initial cell cycle response during axolotl appendage regeneration (7). Although the mechanisms of regenerative regulation through inter-organ communication have been identified, a comprehensive analysis of secreted proteins contributing to tissue regeneration has not yet progressed at the organismal level. *Drosophila* is one of the model organisms that contributes to the systemic aspects of various phenomena through tissue-specific genetic manipulation. Additionally, imaginal discs in *Drosophila* larvae, the epithelial precursors of adult tissues, have been used as models of tissue regeneration (8). In our previous study, we revealed the contribution of the fat body (FB), a counterpart of the mammalian liver and adipose tissue, to wing disc repair (9–11). We found that kynurenic acid (KynA), a humoral metabolite secreted by the FB, was remotely regulated by tachykinin neurons and was necessary for imaginal disc repair (9, 12). Considering that the FB is a major endocrine organ in flies, we have focused on the FB as a source of humoral proteins regulating disc regeneration.

In this study, we attempted to identify secretory proteins that contribute to tissue regeneration. Most proteins are secreted via the ER-Golgi pathway, referred to as the conventional secretory pathway, and undergo glycosylation in the ER lumen or Golgi apparatus (13, 14). Glycosylation is important for the functioning of secreted factors; however, our understanding of its contribution to tissue regeneration is limited. Therefore, we tested whether glycosyltransferases and sugar hydrolases contribute to wing disc regeneration and found that the RNAi knockdown of glycosyltransferases and sugar hydrolases in the FB hampered wing disc repair without any obvious effect on development. Using this finding as a starting point, we conducted a comprehensive search for secreted proteins that contribute to regeneration, using Golgi-derived secretome analyses and RNAi screening.

## Results

### N-glycosylation pathway in the ER and Golgi apparatus contributes to disc regeneration

To induce temporal tissue damage and subsequent wing disc regeneration, a temperature-sensitive diphtheria toxin A domain (DtA^ts^) was used for temporal cell ablation and repair in our previous study (10) (Fig. 1A). Briefly, larvae were reared at 29 °C to stop cell ablation and temporally reared at 18 °C to induce cell ablation. After hatching, we examined the regenerating ability by observing the adult wing phenotype. To manipulate Gal4/UAS-mediated gene expression in the FB, DtA^ts^ was independently induced by the Q system, another binary system, in the wing pouch (WP) region of the wing disc, which becomes an adult wing through metamorphosis. We also examined the effects of genetic manipulation on development in the FB by maintaining the temperature at 29 °C. Using this system, we attempted to identify regenerative regulatory proteins secreted from the FB.

**Figure 1.**
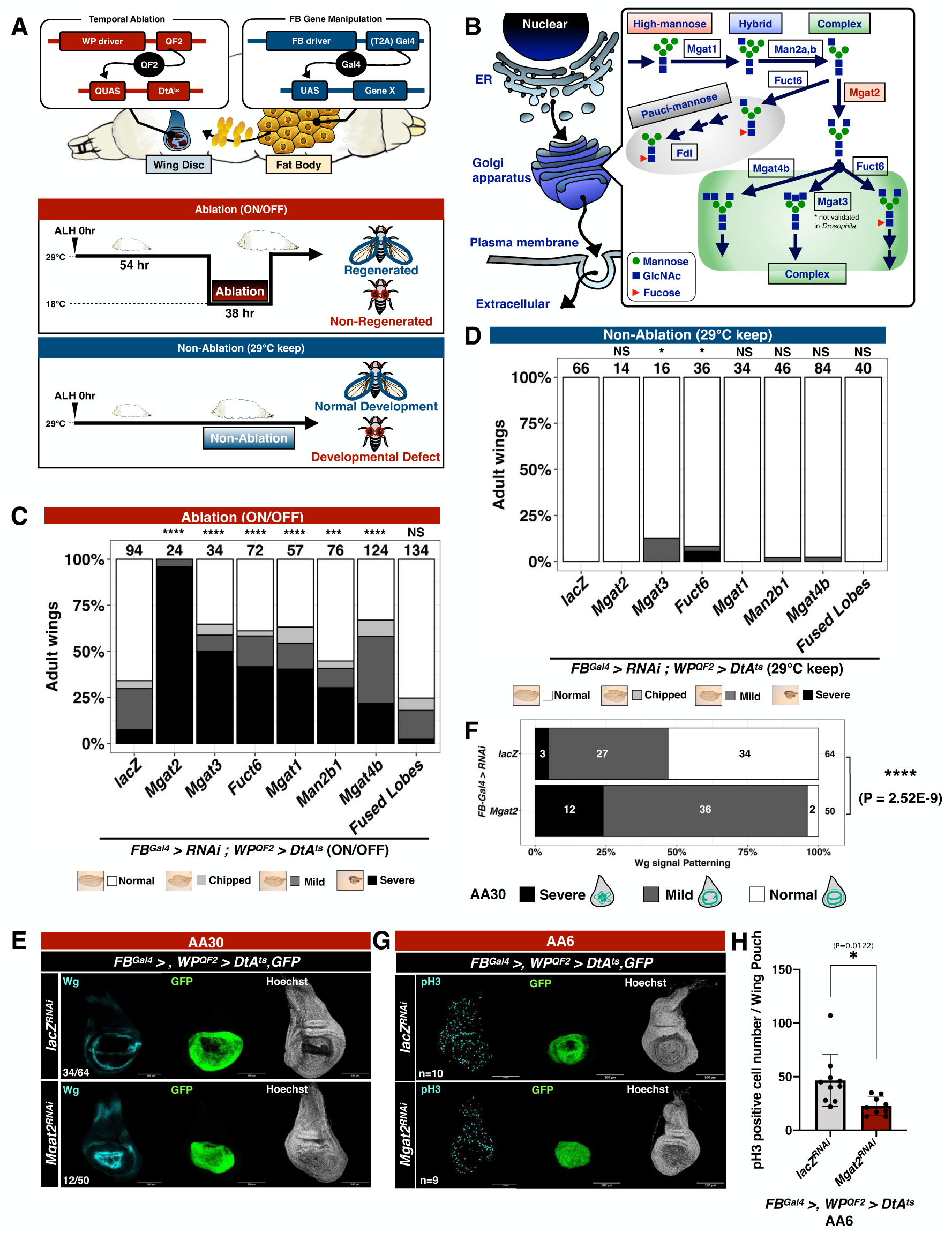
Glycosyltransferases and sugar hydrolases in Fat Body for disc regeneration. (A) Schematic view of temporal disc ablation and gene manipulation in FB. Temporal disc damage is induced by DtA^ts^ at a low temperature (18 °C) for 38 hr and returned to 29 °C after ablation. This condition is indicated as “Ablation”. In a non-damaged condition, temperature shift is not induced and kept at 29 °C to avoid damage induction by DtA^ts^. This condition is indicated as “Non Ablation.” (B) N-glycosylation pathway in the Golgi apparatus. It is important to note that Mgat1, 2, and 4 were functionally validated in *Drosophila*, but Mgat3 was not. (C and D) Comparison of adult wing sizes between ablation (C) and non-ablation (D). RNAi knockdown of N-glycosylation enzymes hampered disc regeneration. RNAi knockdown was induced in FB by *FB^Gal4^*. Statistical analysis was conducted using Fisher’s exact test to compare control (*WP^QF2^>DtA^ts^ ; FB^Gal4^>lacZ^RNAi^*) with treated larvae. NS: not significant, **: p<0.01, ****: p<0.0001. Number of flies is listed above each genotype in the bar graph. (E and F) Representative examples of wing discs developed within the indicated time course. Discs were dissected at 30 hr After Ablation (AA30). FB specific RNAi knockdown of *Mgat2* affected Wg patterning in disc regeneration. Discs were stained with anti-Wg antibody and Hoechst33342. White scale bar, 100 μ m. Abnormality of Wg patterning in wing pouch was classified into 3 groups, and classification was quantified in (F). Fisher’s exact test was applied. ****: p<0.0001. Number of wing discs is listed in image or next to each genotype in the bar chart. (G and H) Representative examples of wing discs developed within indicated time course. Discs were dissected at 6 hr After Ablation (AA6). FB specific RNAi knockdown of *Mgat2* affected cell proliferation in disc regeneration. Discs were stained with anti-pH3 antibody and Hoechst33342. White scale bar, 100 μ m. pH3-positive cell numbers in wing pouch region were quantified in (H). Error bars indicate standard error of the mean. An unpaired t-test was applied. *: p<0.05. Number of wing discs is listed in (G).

First, we examined the effects of general secretion from the FB on disc regeneration. A previous study showed that factors involved in the protein transport pathway contribute to general secretion from the FB(15). RNAi knockdown of several factors involved in the secretory pathway, such as *Rab1*, *Syx18,* and *Rab11*, and endocytosis, such as *AP2α* and *Shibire*, caused a lethal phenotype, indicating their essential role in the FB during development (Fig. S1A, B, C). In contrast, RNAi knockdown of other factors involved in secretion and endocytosis, such as *Sec23*, *deltaCop*, and *Tweek*, increased the percentage of the worsened wing phenotype (Fig. S1B). RNAi knockdown of *Sec23*, *deltaCOP*, and *Tweek*, which causes regeneration failure, had little effect on normal wing formation (Fig. S1C). These results suggest that protein secretion from the FB to wing discs during ablation supports wing disc regeneration, whereas impairment of several protein transport regulators causes lethality.

When humoral proteins are secreted via the conventional secretory pathway, most of them undergo a series of glycosylation steps as they move from the ER to the Golgi apparatus (14). Focusing on the glycosylation process as a characteristic of secretion, rather than the secretory pathway itself, could facilitate screening with a minimal impact on development. Therefore, we investigated whether glycosyltransferases and sugar hydrolases contributed to disc regeneration. N-glycosylation in *Drosophila* is classified into high-mannose, pauci-mannose, complex, and hybrid types. We examined the roles of the enzymes before and after branching of the N-glycosylation pathway (Fig. 1B). We observed that FB-specific RNAi knockdown of the enzyme, *Fused Lobes* (*Fdl*), specifically contributing to the pauci-mannose pathway, did not affect disc repair. However, FB-specific RNAi knockdown of enzymes contributing complex and hybrid N-glycosylation pathway resulted in a worsened wing phenotype after disc damage (Fig. 1C). Furthermore, RNAi knockdown of ER glycosyltransferases *Alg3* and *Alg9,* and sugar hydrolases *GCS1*, and the early phase of glycosyl modification in the Golgi, *Man1b,* and *Man1c*, caused regenerative inhibition (Fig. S1D, E). As expected, FB-specific RNAi knockdown of glycosyltransferases and sugar hydrolases during normal development had no significant effect on wing formation (Fig. 1D and Fig. S1F). Among the glycosyl-modifying enzymes examined, we focused on Mgat2 as it acts on the complex N-glycosylation branch and significantly affects disc-regeneration phenotype (Fig. 1B, C). We confirmed the knockdown efficiency of *Mgat2* (Fig. S2A) by RNAi knockdown of *Mgat2* in the FB with FB Gal4 drives. Although the RNAi knockdown efficiencies of *Mgat1* and *Mgat2* were found to be similar (Fig. S2A, B), the inhibitory effect on regeneration was less pronounced with the RNAi knockdown of *Mgat1* (Fig. 1C). Even after *Mgat1* knockdown, a small amount of *Mgat1* remained (Fig. S2B). Since *Drosophila* have an abundance of high-mannose type glycans (16), even a small amount of *Mgat1* may be sufficient to produce glycans for the next modification. Therefore, the inhibitory effect of the RNAi knockdown of *Mgat1* on regeneration may be limited. Ablation by DtA^ts^ caused a developmental delay, as previously described (10), but RNAi knockdown of *Mgat2* did not affect developmental speed (Fig. S2E), indicating that Mgat2 regulates disc repair by affecting repair processes. Gene expression of glycosyltransferases and sugar hydrolases, except for *Mgat4b*, at 0 hr after injury (after ablation at 0 hr, AA0) and 6 hr (AA6), showed no significant change between ablation and non-ablation conditions (Fig. S3A–E).

Next, we investigated whether Mgat2 in the FB contributed to wing disc remodeling after injury by observing the expression of the morphogen wingless (Wg) and GFP-labeled WP regions (*WP^QF2^>mCD8-GFP*). RNAi knockdown of *Mgat2* in the FB did not significantly affect the WP region area recovery but affected Wg re-patterning at 30 hr after injury (AA30), a late stage of repair (Fig. 1E, F, S2F). This Wg patterning abnormality was not due to excessive cell death by staining for cleaved Dcp1 (cDcp1), an indicator for activated caspase (Fig. S2G, H). Although the recovery area was not affected at AA30, the number of mitotic cells (phospho-histone 3 (pH3)-positive cells) in the early stage of regeneration (AA6) was reduced upon the RNAi knockdown of *Mgat2* in the FB (Fig. 1G, H). Consequently, Mgat2 in the FB is required to restore cell proliferation during the early stages of repair.

### Constructing a system for biotin labeling in the Golgi apparatus

To identify regenerative proteins secreted from the FB, we aimed to comprehensively detect secretory proteins using TurboID, a tool that biotinylates proximal proteins within a 10 nm range (17). In a previous study, BirA*G3, an earlier version of TurboID, was expressed in the *Drosophila* ER lumen in a tissue-specific manner, enabling the detection of secretory proteins in the specific tissues of other organs (2). In our study, we focused on the Golgi apparatus because complex N-glycosylation was found to affect disc regeneration (Fig. 1B–D). We analyzed tissue-specific secretory proteins by fusing TurboID with a glycosyltransferase present in the Golgi apparatus. Given that a remarkable regeneration failure phenotype was observed with FB-specific RNAi knockdown of *Mgat2* (Fig. 1C), we generated *Mgat2-V5-TurboID* fly by fusing C-terminus of Mgat2 with a V5 tag and TurboID (Fig. 2A, B).

**Figure 2.**
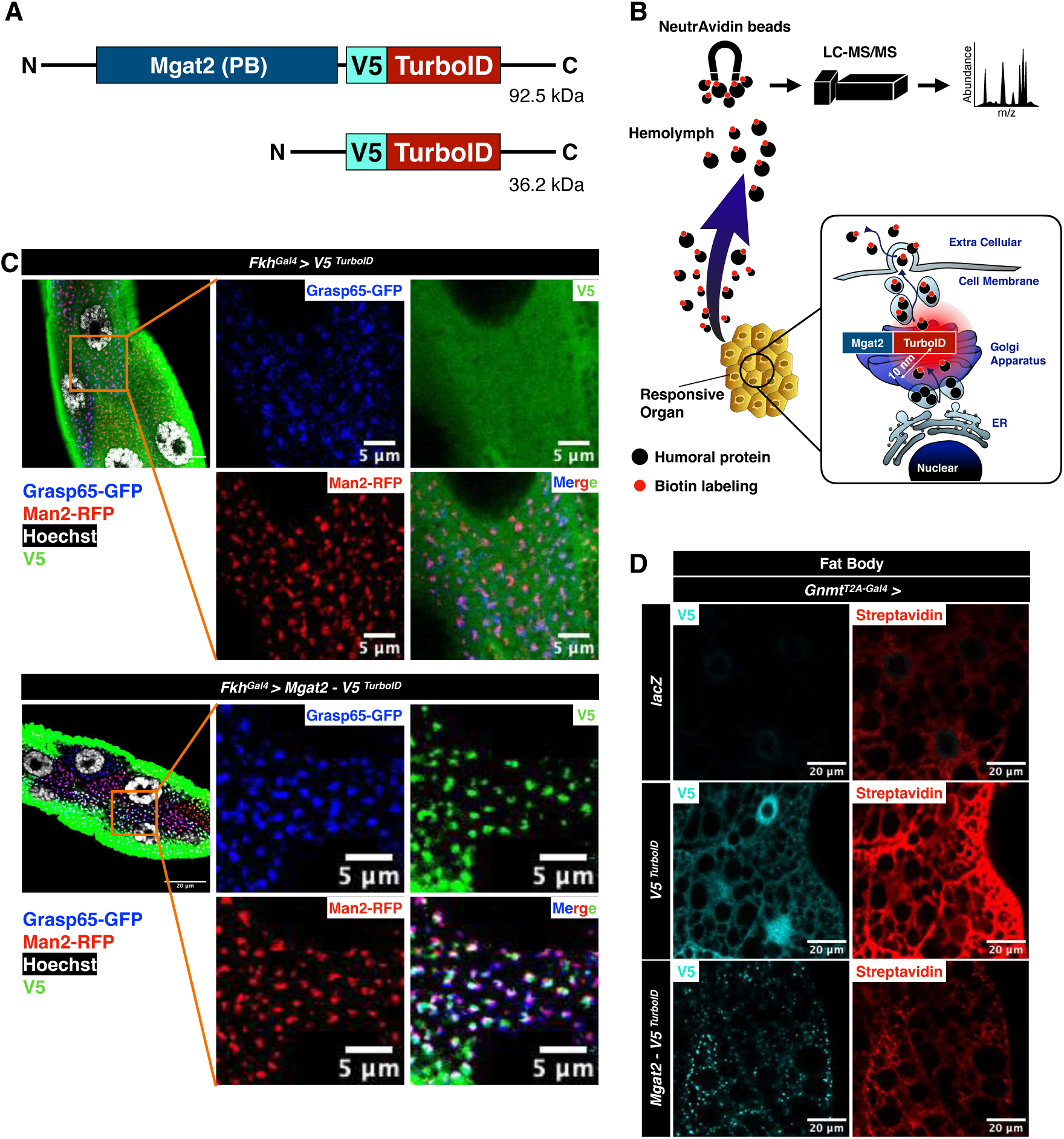
Subcellular localization and biotin-labeling of Mgat2-TurboID. (A) Diagram of V5-TurboID and Mgat2-V5-TurboID (isoform B, PB) constructs. (B) Schematic view of secretome analysis. Tissue-specific expression of *Mgat2-TurboID* enables comprehensive biotin-labeling of secretory pathway proteins via near-labeling in the Golgi apparatus. (C) Cellular localization of Mgat2-V5-TurboID and V5-TurboID. Over-expression of each genotype in salivary gland with *Fkh^Gal4^*. V5 signal (Green) shows localization of each construct. Grasp65 (Blue) is a marker of Cis-Golgi, and Man2 (Red) is a marker of Medial-Golgi. White scale bar, 5 or 20 μ m. Number of samples was 3 for *Fkh^Gal4^>V5^TurboID^*and 4 for *Fkh^Gal4^>Mgat2-V5^TurboID^*. (D) Subcellular localization and biotin-labeling of Mgat2-V5-TurboID and V5-TurboID. Over-expression of each genotype in FB with *Gnmt^T2A-Gal4^*. V5 signal (Cyan) and biotin-labeling (Red) were stained with anti-V5 antibody and Streptavidin-Cy3, respectively. White scale bar, 20 μ m. Number of samples was n=10 in *Gnmt^T2A-Gal4^>lacZ*, n=10 in *Gnmt^T2A-Gal4^>V5^TurboID^* and n=6 in *Gnmt^T2A-Gal4^*>Mgat2-V5*^TurboID^*.

When V5-TurboID was expressed in the salivary gland using *Fkh^Gal4^*for observation, the V5 signal diffused within the cell, whereas the Golgi membrane markers Grasp65-GFP (Cis-Golgi) and Man2-RFP (Medial Golgi) formed puncta (Fig. 2C) (18). In contrast, the expression of Mgat2-V5-TurboID (predicted to localize to the Medial Golgi (19)) resulted in puncta formation, and the puncta of Mgat2-V5-TurboID nearly co-localized in close proximity to Golgi membrane markers (Fig. 2C). Furthermore, we investigated the signal of biotin-labeled proteins and V5 tags in the FB. Unlike lacZ (control), V5-TurboID was diffusely localized to the cytoplasm and nucleus (Fig. 2D). Biotin-labeled proteins marked with streptavidin are localized mainly in the cytoplasm. For Mgat2-V5-TurboID, punctate signals of V5 and biotin-labeled proteins were detected in the cytoplasm (Fig. 2D). These data indicate that Mgat2-V5-TurboID is present in the Golgi apparatus and can label nearby proteins in the Golgi apparatus.

### Analysis of secreted proteins from the FB using Mgat2-TurboID identified a group of proteins responsible for regeneration

To validate whether Mgat2-TurboID (same as Mgat2-V5-TurboID) can label secretory proteins, we examined Mgat2-TurboID expression in the FB. In order to select suitable FB-specific Gal4 drivers for both TurboID and Mgat2-TurboID expression, we examined three FB-Gal4 drivers (FB^Gal4^, Gnmt^T2A-Gal4^, and r4^Gal4^). Both TurboID and Mgat2-TurboID in the FB were observed with Gnmt^T2A-Gal4^ and r4^Gal4^ but not with FB^Gal4^ (Fig. S4A). Consistently, biotin-labeled proteins, detected by streptavidin in both the hemolymph and FB, were present in Gnmt^T2A-Gal4^ and r4^Gal4^ samples but absent in those from FB^Gal4^ (Fig. S4A, B). These results therefore indicate that both TurboID and Mgat2-TurboID can be effectively expressed using Gnmt^T2A-Gal4^ and r4^Gal4^, although TurboID can only be expressed using FB^Gal4^. Given these findings and the technical difficulty of genetic recombination of *r4^Gal4^* and *WP^QF2^*, we selected Gnmt^T2A-Gal4^ for subsequent Mgat2-TurboID analyses. Western blot analysis of biotinylated proteins in disc regeneration showed that *Drosophila* wandering 3^rd^ instar larvae expressing Mgat2-TurboID exhibited labeled proteins in both hemolymph and the FB (Fig. 3A, B). In the hemolymph, labeled Mgat2-TurboID proteins were detected at much higher levels compared to lacZ and TurboID, and the banding patterns of Mgat2-TurboID closely resembled those of BirA*G3-ER (Fig. S4C). Cytosolic αTubulin was not detected in the hemolymph samples. In FB, TurboID strongly labeled cellular proteins (Fig. S4D). Additionally, Mgat2-TurboID and BirA*G3-ER showed a similar banding pattern in the hemolymph and FB, implying that Mgat2-TurboID efficiently labeled the proteins of secretory pathway (Fig. S4C, D).

**Figure 3.**
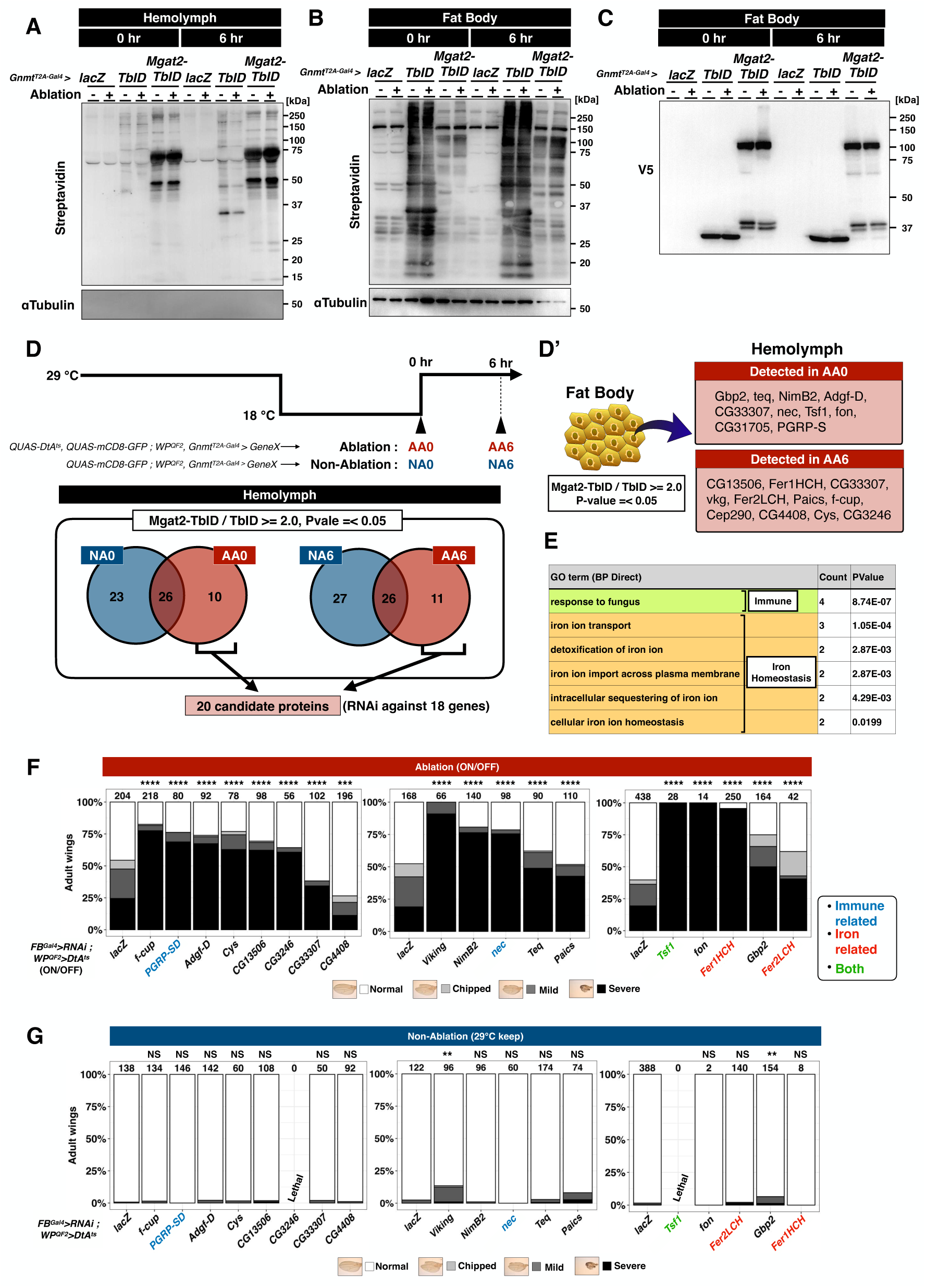
Secretome analysis identified Fat Body-derived iron- and immune-related proteins are necessary for disc regeneration. (A and B) Western blotting for biotinylated proteins (detected by Streptavidin-HRP) in hemolymph (A) and FB (B). *LacZ* (Control without TurboID), *TurboID* (TbID), and *Mgat2-TurboID* (Mgat2-TbID) were expressed in FB with *Gnmt^T2A-Gal4^*. Samples were collected both in Ablation and Non-Ablation conditions and at 0 and 6 hr time points. αTubulin was used as an internal control. 10–20 larvae were used for each lane. (C) Western blotting for V5-tagged proteins in FB. V5-TurboID and Mgat2-V5-TurboID were estimated to be 36.2 kDa and 92.5 kDa, respectively. 10–20 larvae were used for each lane. (D, D’) Scheme for temperature change and sampling. Analysis of biotinylated proteins in hemolymph and FB at two time points. Under each condition, proteins detected with *Mgat2-TurboID* at more than twice the amount detected with *V5-TurboID* (p < 0.05) were extracted. CG33307 in hemolymph overlapped in both AA0 and AA6. 15–25 larvae were used for single samples, and biological replicates were n=3. (E) GO analysis of 20 proteins in (D) with DAVID. (F and G) RNAi screening of candidate proteins in Ablation (F) and Non-Ablation (G). Immune-related proteins are indicated in blue, iron-related proteins are indicated in orange, and proteins involved in both are indicated in magenta. Statistical analysis was conducted using Fisher’s exact test to compare control (*WP^QF2^>DtA^ts^ ; FB^Gal4^>lacZ^RNAi^*) with treated larvae. NS: not significant, **: p<0.01, ***: p<0.001, ****: p<0.0001. Number of flies is listed above each genotype in the bar graph.

Although Mgat2-TurboID appeared to efficiently label the proteins within the secretory pathway, membrane-anchored Golgi glycosyltransferase was cleaved at the N-terminus, and catalytic enzymes were secreted into the extracellular space (20). In order to determine whether TurboID or Mgat2-TurboID itself is secreted, we investigated TurboID and Mgat2-TurboID secretion by expressing them in the FB using *Gnmt^T2A-Gal4^* and *r4^Gal4^*, and found their presence in the hemolymph (Fig. S4B). However, since TurboID functions in ATP-dependent manner and extracellular ATP levels are extremely low in the human plasma (21), it can be speculated that TurboID does not function well in the hemolymph. When taken together, these observations support the conclusion that TurboID and Mgat2-TurboID act as biotinylating enzymes within the FB and that the observed protein labeling occurs intracellularly prior to secretion.

Additionally, we have expressed *TurboID* and *Mgat2-TurboID* in several cell types and tissues using *nSyb^Gal4^*for neurons, *Pros^Gal4^*for neural cells and gut enteroendocrine cells, *Mef2^Gal4^*for muscle, *Pxn^Gal4^* for hemocytes, and *Fkh^Gal4^*for the salivary glands (Fig. S4B, E, F). Labeled proteins were detected in the hemolymph by expressing *TurboID* and *Mgat2-TurboID* using *Mef2^Gal4^* and *Pxn^Gal4^*. Therefore, *TurboID* and *Mgat2-TurboID* can be used in several cells and tissues other than FB.

To identify proteins specifically secreted from the FB during regeneration, we conducted FB-specific protein labeling and proteomics analysis of the hemolymph and FB at the early stage of disc repair (AA0 and AA6) (Fig. 3D). Western blot analysis indicated no significant difference in labeled proteins detected in the hemolymph or FB at any time point between the ablation and non-ablation groups (Fig. 3A, B). There were also no differences in the amounts of TurboID (TbID) or Mgat2-TurboID (Mgat2-TbID) proteins present in the FB, regardless of the time point or ablation (Fig. 3C). Mgat2-TurboID and BirA*G3-ER showed similarly labeled proteins in FB proteomics, as seen in western blot analysis (Fig. S5A). In the hemolymph samples, Mgat2-TurboID and BirA*G3-ER shared many common factors under all conditions (Fig. S5B). Cellular component and pathway analyses were performed using the Database for Annotation, Visualization and Integrated Discovery (DAVID) (https://david.ncifcrf.gov). Cellular component analysis indicated that most of the proteins labeled by V5-TurboID were present in the cytosol and nucleus under all conditions (Table S1), consistent with the observation that V5-TurboID expression was also observed in the cytosol and nucleus (Fig. 2C, D). Membrane proteins were the top hits in both Mgat2-TurboID and BirA*G3-ER but not in TurboID in the FB (Table S1). ER-related proteins were also more enriched in both BirA*G3-ER and Mgat2-TurboID, and proteins related to the Golgi apparatus were enriched in Mgat2-TurboID under all conditions (Table S1). Furthermore, well-known major secreted proteins from the FB, such as Lsp1α,γ, β, and Lsp2, were labeled more in the Mgat2-TurboID samples when compared to those of TurboID samples in the hemolymph (Fig. S5C). These data indicate that secretory pathway proteins were more specifically labeled by Mgat2-TurboID and BirA*G3-ER than by TurboID. In the pathway analysis, N-glycosylation-related terms were more frequently detected in Mgat2-TurboID than in others, whereas terms related to ER function were enriched in both BirA*G3-ER and Mgat2-TurboID (Table S2). In summary, these results suggest that the proteins detected in both Mgat2-TurboID and BirA*G3-ER were relatively similar, whereas Mgat2-TurboID more specifically labeled secretory proteins within the Golgi apparatus.

To identify proteins secreted from the FB into the hemolymph during regeneration, we compared the labeled proteins, which were two times higher in Mgat2-TurboID than in TurboID (Fig. 3D, D’, and Table S3). We identified 20 candidate regenerative secretory proteins from the FB in AA0 and AA6 by focusing on the factors that were undetected in non-ablation (NA) but detected only in after-ablation (AA). GO analysis of these regenerative candidate proteins using DAVID revealed the upregulation of factors related to innate immunity and iron homeostasis (Fig. 3E). In addition, we compared the labeled proteins, which were two times higher in BirA*G3-ER than in TurboID (Fig. S5D, D’, and Table S3). We identified 14 proteins as regenerative candidate proteins by focusing on the factors that were undetected in NA but detected only in AA. GO analysis of these proteins also revealed the upregulation of factors related to iron homeostasis and innate immunity (Fig. S5E). These results suggest that the factors contributing to the regeneration of wing discs are related to iron binding and innate immunity.

To confirm the importance of FB-derived candidate regenerative proteins in disc regeneration, we conducted RNAi screening of 18 genes among 20 candidates (Fig. 3F, G). Most of selected factors were required for disc regeneration. FB-specific RNAi knockdown of iron or immune-related factors, *transferrin 1* (*tsf1*), *Fer1HCH*, *Fer2LCH*, *PGRP-SD*, and *necrotic* (*nec*), had severe negative effects on disc regeneration (Fig. 3F, G). These data indicate a non-autonomous requirement for innate immunity and iron-related factors for disc regeneration.

## Discussion

In this study, we found that FB glycosylation was non-autonomously required for disc repair (Fig. 1C). To determine the humoral factors involved in disc repair, we established Mgat2-TurboID, a Golgi-specific labeling tool. This system enabled us to perform secretome analysis involving the Golgi apparatus and identify secretory proteins crucial for disc regeneration, including innate immune regulators and iron mediators (Fig. 3D–F). The contribution of immunity to tissue damage response and inflammatory regulation in tissue repair has been reported and discussed across species, such as mammals and flies (22, 23). In flies, hemocytes, one of the immune regulatory cells, are recruited to damaged wing discs and subsequently activate the JAK/STAT pathway in the hemocytes and FB (24). However, it remains unclear how the Toll and Imd pathway-associated factors detected in the hemolymph proteomics play their respective roles. Recent *Drosophila* studies have suggested a link between iron and innate immunity. During septic injury, hemolymph iron is sequestered into the FB, a phenomenon known as nutritional immunity (25). Iron uptake during infection in the FB suppresses the growth of pathogenic bacteria. During this defense response, the iron transporter Tsf1 is secreted into the hemolymph to sequester serum iron, depending on the immune signaling of the Toll and Imd pathways. In our study, we found that innate immune factors, such as PGRP-SD, nec, and the iron transporter Tsf1, contributed to disc repair (Fig. 3F). Tsf1 is transcriptionally regulated by Toll and Imd signaling (25), and nec is an inhibitor of Toll pathway (26), and PGRP-SD is an activator of Imd pathways (27, 28). These findings suggest that iron metabolism may be regulated by innate immune pathway. Further investigation is essential to elucidate the role and relationship between iron mediators and immune signaling in disc regeneration.

As previously mentioned, transferrin (Tsf) is an iron transport protein, and tsf1 is one of the candidate proteins in this disc regeneration study. In insects, Tsf1 can bind ferric iron, and ferritin can store approximately 3,000 ferric ions in the ferritin shell. They transport or uptake iron from various organs (29, 30). In mammals, iron uptake and release by blood cells play a role in tissue injury (31, 32). In injured cell populations, damage mitigation and repair responses via iron regulation by blood cells proceed locally to the injury site. In the early stages of injury, macrophages protect themselves from harmful iron released from the injured site and limit bacterial growth by absorbing and sequestering iron at the injury site. Such iron sequestration leads to anemia of inflammation observed in chronic inflammatory diseases, such as chronic infections, autoimmune diseases, cancer, and chronic kidney disease (31, 32). Subsequently, during the repair phase, macrophages release iron around the site of repair, stimulating cell proliferation and angiogenesis at the injury site (32). Thus, in mammalian tissue repair, the process begins with iron sequestration by macrophages, followed by the subsequent iron supply. Kupffer cells in the liver, like macrophages, are major transferrin producers and are central to systemic iron metabolism (32). However, it has not been shown how iron metabolism regulation by the liver affects remote tissue repair. In this study, we found that iron metabolism regulators generated through secretory pathway in the FB remotely contribute to tissue repair (Fig. 3). How FB-derived iron regulators affect hemocytes remains unclear and warrants further investigation.

We have reported that kynurenine (Kyn) metabolism in the FB is enhanced in response to tissue injury and that KynA secreted from the FB remotely supports tissue repair (9, 12). Vermilion (mammalian tryptophan 2,3-dioxygenase), the rate-limiting enzyme for Kyn production, is an iron-containing oxygenase (33). Therefore, iron regulation in the FB may affect the activity of heme-containing enzymes such as Vermilion and may contribute to remote tissue repair by regulating iron metabolism in the FB.

Previous studies have examined the role of glycosylation in tissue regeneration, but glycosylation does not always promote regeneration. For instance, knock-out of N-glycosylation hinders axon regeneration in *Caenorhabditis elegans* (34). In contrast, in zebrafish hair cell regeneration, the repair process is enhanced by deficiency of Mgat5, an N-glycosylation enzyme (35). In mouse heart regeneration, a deficiency in N-glycosylation of hFSTL1 promotes regeneration (36). As shown above, glycosylation shows different requirements in tissue repair depending on factors, conditions, species, and tissues. In this study, we attempted to comprehensively search for the FB-derived secretory proteins that influence disc regeneration using Mgat2-TurboID (Fig. 3D, E). Mgat2-TurboID allowed us to identify humoral factors secreted via the Golgi apparatus. Glycosylation affects not only secretion but also protein structure and activity. In addition to a comprehensive search for secreted proteins, a deeper understanding of regenerative factors will require further investigation into glycosylation, including glycoproteomic approaches, such as pGlyco3 (37).

The FB serves as a major endocrine organ, comparable to the liver and adipose tissue in mammals, and is responsible for most protein production through the secretory pathway (38). In this study, we employed a proximity-dependent secretory protein labeling method to investigate the secretory pathway proteins during disc regeneration. In previous studies, labeling enzymes were fused to the ER lumen to create proximity-dependent secretory protein labeling tools, such as ER-BioID(39), BirA*G3-ER(2), and ER-TurboID(40–43). Using these tools, most secretory proteins from specific organs have been identified in the hemolymph or blood. We fused the labeling enzyme with a glycosyltransferase in the Golgi apparatus to create Mgat2-TurboID, as our initial focus was on complex-type N-glycosylation in disc regeneration. As a tool for detecting secretory proteins, there was a large overlap in the FB-derived labeled humoral proteins between Mgat2-TurboID and BirA*G3-ER during disc regeneration (Fig. S5A, B). However, we observed that Mgat2-TurboID labeled Golgi proteins more efficiently than BirA*G3-ER. Furthermore, a higher number of membrane proteins was detected in Mgat2-TurboID than in BirA*G3-ER (Table S1). Considering these observations, our Golgi-specific labeling platform enhances the specificity of proximity labeling within the secretory pathway and is widely applicable for future studies on inter-organ protein trafficking and membrane protein labeling.

## Experimental Procedures

### Fly strains and genetics

All flies were reared on Saf-instant and oriental corn meal medium (hereafter Saf or oriental food) at 25 °C. The detailed food composition per liter of Saf and Oriental foods is shown in Tables 1 and 2. The fly lines and genotypes used in this study are presented in Tables S4 and S5.

**Table 1.** Saf-instant fly food composition per liter.

| Ingredient | Amount | Brand and catalog number (if applicable) |
| --- | --- | --- |
| Agar | 8 g | KISHIDA CHEMICAL, 260-01705 |
| Glucose | 60 g | WAKO, 049-31165 |
| Dry yeast | 60 g | Saf-instant RED |
| Corn flour | 40 g | NIPPON, 100016 |
| Propionic acid | 3 mL | WAKO 163-04726 |
| 10% butyl p-hydroxybenzoate | 1.5 mL | WAKO 132-02635 |

**Table 2.** Oriental fly food composition per liter.

| Ingredient | Amount | Brand and catalog number (if applicable) |
| --- | --- | --- |
| Agar | 8 g | KISHIDA CHEMICAL, 260-01705 |
| Glucose | 100 g | WAKO, 049-31165 |
| Dry yeast | 40 g | Oriental |
| Corn flour | 40 g | NIPPON, 100016 |
| Propionic acid | 3 mL | WAKO 163-04726 |
| 10% butyl p-hydroxybenzoate | 5 mL | WAKO 132-02635 |

### Temperature shift protocol for temporal ablation

Embryos were laid at 25 °C for 4 hr, and we defined this time point as after egg laying (AEL) as 0 h. The temperature was maintained at 25 °C and shifted to 29 °C, 24 h after AEL 0 hr. For ablation with *WP^QF2^>DtA^ts^*, the temperature was raised to 29 °C for 54 hr, and shifted to 18 °C from the early third instar larval stage. After maintaining an 18 °C for 38 hr, the temperature was returned to 29 °C and flies were either allowed to pupate and eclose, or were dissected at the time points indicated as 0, 6, or 30 hr After Ablation (AA). For proteomics analyses, developmental speed measurements, and qPCR analysis, larvae without DtA^ts^ were used as the non-damaged control, and samples were collected at 0 or 6 h non-ablation (NA), at the same time points as AA.

### Wing Size Assessment

The flies were preserved in a mixture of ethanol/glycerin (3:1). We observed wing phenotypes by microscopy and manually classified the wing size into four categories: 1) normal: intact wings; 2) chipped: chipped or crumpled wings having a size >90% of control; 3) mild: wings with intermediate phenotypes, or between ‘Chipped’ and ‘Severe’ groups, 4) severe: wings measuring <30% of normal size. When counting the wing regeneration rate, we noted the statistical analysis values and number of samples (N) above each genotype bar. To take wing pictures, we dehydrated flies in 100% ethanol for 4-24 hr, following dissection of wings in 100% ethanol and mounting them in Euparal mounting medium (WALDECK GmbH & Co KG, DIVISION CHROMA, 35358-86). Images were captured with a Leica microscope DM5000.

### Quantification of developmental speed

Time to pupation was measured to determine the developmental speed. The injured and uninjured larvae were subjected to temperature changes according to temperature shift protocol, and the number of pupated flies was subsequently counted at 10 time points of AEL at 96, 105, 120, 129, 144, 153, 168, 177, 192, and 201 hr. Pupae number at AEL of 201 hr was set to 1.0, and the developmental speed from larvae to pupae was measured as the pupal fraction at each time point.

### Immunohistochemistry

Discs, fat bodies, and salivary glands were dissected in PBS and fixed with a 4% paraformaldehyde PBS solution for 20 min. Discs were washed with 0.1% Triton X-100 PBS solution (PBST). Fat bodies and salivary glands were washed with 0.3% Triton X-100 PBST. The antibodies used were as follows: rabbit anti-cleaved Dcp1 (Asp216) 1:200 (Cell Signaling, 9578S), mouse anti-Wg (4D4) 1:500 (Developmental Studies Hybridoma Bank), rat anti-phospho-histone H3 (Phospho S28) 1:500 (Abcam, ab10543), rabbit anti-RFP 1:500 (Abcam, ab62341), and mouse anti-V5 1:500 (Invitrogen, R96025). The secondary antibodies used in this study were as follows: Alexa Fluor antibodies (Thermo Fisher Scientific, A-21202, A-21206, A-21208, A-31571, A-31573, A-48272) and AffiniPure antibodies (Jackson ImmunoResearch, 715-165-150, 711-165-152, 712-166-153) diluted at 1:500. The probes used were as follows: Hoechst 33342 (H3570) and Rhodamine Phalloidin (R415) diluted 1:200 (Life Technologies) and Streptavidin-Cy3 (Jackson ImmunoResearch, 016-160-084) diluted 1:500. Confocal images were captured using a Leica SP8 microscope. Quantification of pH3-positive cells and cell death in the WP region was performed using Fiji software. WP region is the area of the wing pouch labeled with WP^QF2^ and QUAS-mCD8-GFP in the wing disc.

### Fly RNA extraction and cDNA synthesis

RNA extraction was performed using the RNeasy Micro Kit (QIAGEN) and the ReliaPrep^TM^ RNA Tissue Miniprep System (Promega) according to the manufacturer’s instructions. RT-qPCR was performed according to the protocol for nonfibrous tissues using the ReliaPrep^TM^ RNA Tissue Miniprep System (Promega). Briefly, the fat bodies of three third-instar larvae were dissected in PBS and homogenized in 1.5 mL tubes containing lysis buffer. Then, the samples were stored at -80 °C according to the protocol of ReliaPrep^TM^ RNA Tissue Miniprep System (Promega). cDNA was synthesized from 400 ng of total RNA using PrimeScript RT Master Mix (Perfect Real Time) (Takara Bio).

### Expression constructs

cDNA samples for construction were prepared according to the RNeasy Micro Kit protocol (QIAGEN, 74004) and PrimeScript RT Master Mix (Perfect Real Time) (Takara Bio, RR036A). In QIAGEN protocol, six adults or six *UAS-lacZ^RNAi^* larvae were homogenized in 150 μ L of QIAzol, followed by the addition of 350 μ L QIAzol and incubation for 30 minutes. Subsequently, samples were stored at -80 °C according to the protocol of RNeasy Micro Kit (QIAGEN, 74004). For constructing *UAS-Mgat2-V5-TurboID*, the isoform RB of *Mgat2* was cloned from the cDNA extracted from adult or third instar larvae of the *UAS-lacZ^RNAi^* using

forward primer 5′-

AAATCAAAGGATCCCCAAAATGTCGAAAATG

and the reverse primer 5′-

GGGGATGGGCTTGCCCCTCGTGGCCAGCGTC

*UAS-Mgat2-V5-TurboID* and *UAS-V5-TurboID* were constructed using the *pAct5-V5-TurboID* plasmid, which was generously provided by Dr. Shinoda of the University of Tokyo (44). The V5-TurboID fragment for UAS-V5-TurboID was amplified from the *pAct5-V5-TurboID* plasmid, by PCR using the forward primer 5′-

AAATCAAAGGATCCCCAAAATGGGCAAGCCC

and the reverse primer 5′-

TAGTGGTACCCTCGATCACTGCAGCTTTTC

The V5-TurboID fragment for UAS-Mgat2-V5-TurboID was amplified from the *pAct5-V5-TurboID* plasmid, by PCR using the forward primer

5′-GGCAAGCCCATCCCCAACCCC

The reverse primer had the same sequence as that of The V5-TurboID fragment for *UAS-V5-TurboID*. The amplified fragments were inserted into *pUASz1.0* (DGRC, #1431) vector with *Xho*I restriction using the NEBuilder HiFi DNA Assembly Master Mix. This sequence was confirmed by Eurofins, Inc. The vector was injected into the transgenic *y1 w67c23; P{CaryP}attP40* strain. Transgenic flies were generated by Best Gene Inc.

### Quantitative RT-PCR

cDNA was diluted tenfold with Milli-Q, and RT-qPCR was performed using TB Green Premix Ex Taq ^TM^ II (Tli RNaseH Plus) (Takara Bio, RR820L) to a total volume of 10 μ L. Quantitative PCR was performed using the Quantstudio 6 Flex Real-Time PCR system (Thermo Fisher Scientific) according to the standard protocol provided with the software. The primer sequences are as follows:

*RNA pol II* Forward primer 5’- CCTTCAGGAGTACGGCTATCATCT,

*RNA pol II* Reverse primer 5’- CCAGGAAGACCTGAGCATTAATCT ;

*Mgat2* Forward primer 5’- ATGTCGAAAATGAGGGGTCGC,

*Mgat2* Reverse primer 5’- ACTCCGCATGTAGAAGTTGTTG ;

*Fused Lobes* Forward primer 5’- TTCATCCTGACGGTGCTCTAC,

*Fused Lobes* Reverse primer 5’- GTATGGGAAAGCCACTAGCATC ;

*Mgat4b* Forward primer 5’- CAGTTCGTGATTGGAGTACCC,

*Mgat4b* Reverse primer 5’- GGATCAGACAGTCCACGGTT ;

*Mgat3* Forward primer 5’- ATGCAGATGTGAAAGTGGCTG,

*Mgat3* Reverse primer 5’- GTGCTACCGCCTCGTTTACTG.

*Mgat1* Forward primer 5’- CCAAGAACGTGTTTGAGTTCGT,

*Mgat1* Reverse primer 5’- CAGCTCCGCCGATATTTCG.

### Western Blotting

Fat bodies and hemolymph from 10–20 larvae were collected. Fat body was collected in 55 μL of RIPA buffer (1 M Tris-HCl, pH 8.0 50 mL / 5 M NaCl 30 mL / 5 g Sodium Deoxycholate / 1 g SDS / 1 L SDS g / NP-40 10 mL / MQ up to 1 L) containing 1x protease inhibitor (cOmplete™, EDTA-free Protease Inhibitor Cocktail [Roche, 16829900]). After centrifugation at 10,000 × g, 4 °C for 10 min, 45 μL of the intermediate layer was collected and stored at -80 °C. Larval hemolymph was collected in 15 μL ice-cold PBS by pinching the epidermis using forceps, then the extracts were mixed with 30. μL of PBS with 1x protease inhibitor (cOmplete™, EDTA-free Protease Inhibitor Cocktail [Roche, 16829900]). After centrifugation at 1,000 × g, 4 °C for 3 min to remove hemocytes, 40 μL of the supernatant was collected and stored at -80 °C. After protein quantification using the Pierce^TM^ BCA Protein Assay Kit (Thermo Fisher Scientific, 23225), 6 × Laemmli sample buffer (1 M Tris-HCl pH 6.8, 12% SDS, 0.6% bromophenol blue, 15% 2-ME) was added, and samples were boiled at 95 °C for 5 min. Samples were separated by 10 % SDS-PAGE. The proteins were then transferred to Immobilon-P PVDF membranes (Millipore, IPVH00010) at 25 V, 2.5 mA for 20 min using the Trans-Blot Turbo Transfer System (Bio-Rad, 1704150). The membranes for streptavidin blotting were blocked with 3% BSA overnight at 4 °C. And streptavidin-horseradish peroxidase (HRP) was diluted x 1/20,000 in 3% BSA and treated on the membrane for 1–2 h at room temperature. The membranes for antibody staining were blocked with 4% Difco^TM^ skim milk (Becton, Dickinson and Company, 232100)/TBST for 10 min at room temperature. Primary antibodies were diluted in 4% Difco^TM^ skim milk and treated on the membrane overnight at 4 °C. After three washes with TBST at room temperature, the secondary antibody was diluted in 4% skim milk and treated on the membrane for 1–2 h at room temperature. The secondary antibody was removed, washed three times with TBST at room temperature, and then treated with Immobilon Western Chemiluminescent HRP Substrate (Millipore, WBKLS0500). Signals were detected using FUSION SOLO 7S. EDGE (Vilber-Lourmat).

### Purification of biotinylated proteins

Biotin food was prepared by adding 1 mM Biotin (FUJIFILM Wako Pure Chemical Corporation, 132-02635) to saf food to a final concentration of 100 μM. Animals were placed in vials containing biotin-containing food for injury experiments. Samples were extracted as described in the western blotting section. Purification of the labeled proteins was conducted as previously described (44) with some modifications. Briefly, 150 μg of biotinylated protein-containing lysate was subjected to FG-NeutrAvidin beads (Tamagawa Seiki Co., Ltd, TAS8848 N1171) purification. FG-NeutrAvidin beads (25 μL, approximately 500 μg) were washed three times with RIPA buffer. Benzonase (Merck, 70746-4)-treated biotinylated protein samples suspended in 1 mL RIPA buffer were incubated overnight at 4 °C. Conjugated beads were magnetically isolated and washed with 500 μL of ice-cold RIPA buffer solution, 1 M KCl solution, 0.1M Na2CO3 solution, and 4 M urea solution. For LC–MS/MS analysis, the purified samples were washed with 500 μL ultrapure water (FUJIFILM Wako Pure Chemical Corporation, 214-01301) and 500 μL of 50 mM ammonium bicarbonate (Sigma-Aldrich, A6141-25G). The samples were then mixed with 50 μL of 0.1% RapiGest SF (Waters, 186001861) diluted in 50 mM ammonium bicarbonate as anionic surfactant, and 5 μL of 50 mM TCEP (Tris(2-carboxyethyl) phosphine hydrochloride, Sigma, C4706) was subsequently added as a reducing agent. The samples were incubated at 60 °C for 5 min, and then 2.5 μL of 200 mM MMTS (methyl methanethiosulfonate, Thermo Fisher, 23011) was added. One microgram sequence-grade modified trypsin (Promega, V5111) was then added for on-bead trypsin digestion at 37 °C for 16 h. The beads were then magnetically isolated, and 60 μL of the supernatants was collected. Then, 3 μL of 10% TFA (trifluoroacetic acid, FUJIFILM Wako Pure Chemical Corporation, 206-10731) was added to the supernatants, and the samples were incubated at 37 °C for 60 min with gentle agitation. The samples were then centrifuged at 20 000 × g, 4 °C for 10 min. The peptides were desalted and purified using a GL-tip SDB (GL Sciences, 7820-11200) following the manufacturer’s instructions. The samples were speed-vacced at 45 °C for 30 min (Tommy Seiko, CC-105) and dissolved in 25 μL of 0.1% formic acid (Kanto Chemical, 16245-63). The samples were then centrifuged at 20 000 × g, 4 °C for 10 min, and the supernatants were collected. The peptide concentrations were determined using the BCA assay (Thermo Fisher Scientific, 23225). Finally, 250 ng of the purified protein was subjected to LC–MS/MS analysis.

### LC-MS/MS analysis

Samples were loaded onto an Acclaim PepMap 100 C18 column (75 μm × 2 cm, 3 μm particle size and 100 Å pore size; Thermo Fisher Scientific, 164946) and separated using a nano-capillary C18 column (75 μm × 12.5 cm, 3 μm particle size, Nikkyo Technos, NTCC-360/75-3-125) in an EASY-nLC 1200 system (Thermo Fisher Scientific). The elution conditions are listed in Table 3. The separated peptides were analyzed using QExactive (Thermo Fisher Scientific) in the data-dependent MS/MS mode. The parameters for MS/MS analysis are listed in Table 4. The collected data were analyzed using Proteome Discoverer (PD) 2.2 software with the Sequest HT search engine. The parameters for the PD 2.2 analysis are listed in Tables 5 and 6. Peptides were filtered at a false discovery rate of 0.01 using the Percolator node. Label-free quantification was performed based on the intensities of the precursor ions using a precursor-ion quantifier node. Normalization was performed using the total amount of peptides in all average scaling modes. Proteins with 1.5 or 2-fold higher abundance ratios relative to the lacZ or TurboID controls were considered for further analysis. The MS proteomics data were deposited in the ProteomeXchange Consortium via the jPOST partner repository with the dataset identifier PXD054313 for FB samples and PXD055017 for hemolymph samples.

**Table 3.**
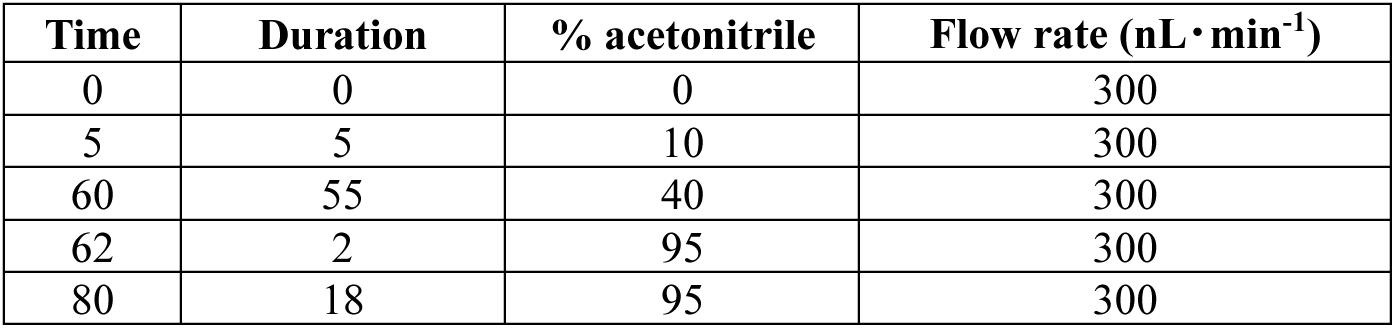
LC settings: Gradient settings for the LC analysis.

| Time | Duration | % acetonitrile | Flow rate (nL · min<sup>-1</sup>) |
| --- | --- | --- | --- |
| 0 | 0 | 0 | 300 |
| 5 | 5 | 10 | 300 |
| 60 | 55 | 40 | 300 |
| 62 | 2 | 95 | 300 |
| 80 | 18 | 95 | 300 |

**Table 4.**
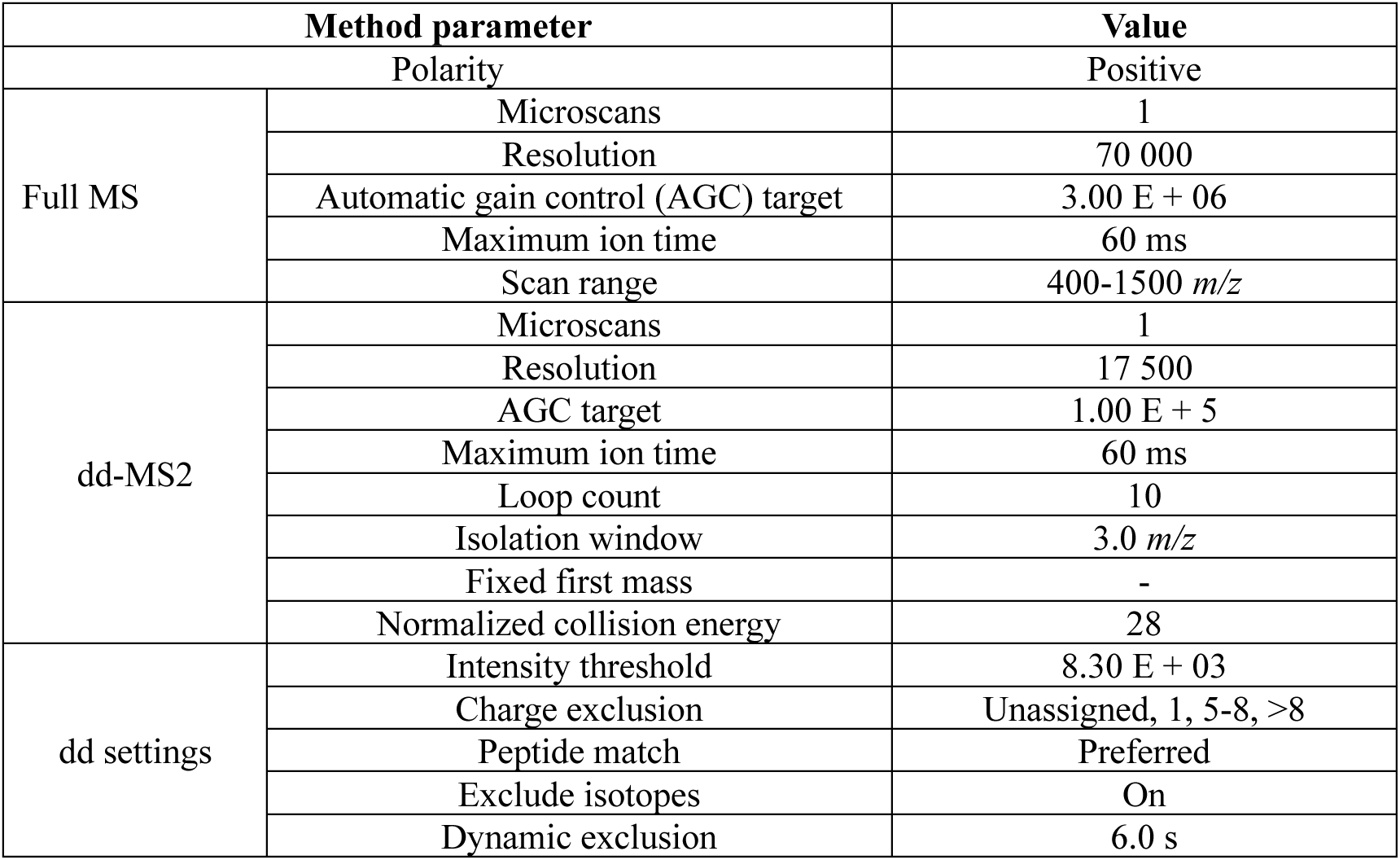
MS/MS settings: Settings used for the MS/MS analysis.

**Table 5.** Proteome Discoverer 2.2 settings: Major Proteome Discoverer 2.2 settings for the processing workflow of proteomics analysis.

| Name of Node | Name of Parameter | Template | This analysis |
| --- | --- | --- | --- |
| Spectrum Files<br>RC | Protein Database | None | Drosophila_melanogaster_uniprot-proteome_UP000000803.fasta |
|  | 1.<br>Static<br>Modification | Carbamidomethyl /<br>+57.021 Da (C) | Methylthio / +45.988 Da (C) |
| Sequest HT | Protein Database | None | Drosophila_melanogaster_uniprot-proteome UP000000803.fasta |
|  | Fragment Mass<br>Tolerance | 0.6 Da | 0.02 Da |
|  | 2.<br>Dynamic<br>Modification | None | GG / +114.043 Da (K) |
|  | 1.<br>Static<br>Modification | Carbamidomethyl /<br>+57.021 Da (C) | Methylthio / +45.988 Da (C) |

**Table 6.** Proteome Discoverer 2.2 settings: Major Proteome Discoverer 2.2 settings for the consensus workflow of proteomics analysis.

| Name of Node | Name of Parameter | Template | This analysis |
| --- | --- | --- | --- |
| MSF Files | Merge Mode | Globally by Search Engine<br>Type | Do Not Merge |
| Precursor Ions<br>Quantifier | Normalization<br>Mode | Total Peptide Amount | None |
|  | Ratio Calculation | Pairwise Ratio Based | Summed Abundance Based |
|  | Hypothesis Test | ANOVA (Background<br>Based) | ANOVA (Individual<br>Proteins) |

### Analysis of secreted regenerative proteins based on proteomics data

For the analysis of proteomics abundance ratio, proteins with Mgat2-TurboID/TurboID higher than 2.0 and a p-value below 0.05 were selected. Candidate proteins detected in the injury condition were extracted. Functional annotation, visualization, and integrated discovery were performed using the web tool DAVID (https://david.ncifcrf.gov/).

### Statistical analysis

Data are presented as stacked bar graphs, bar charts, or mean ± standard error. To analyze the independence of the cross-tabulation tables used in the stacked bar graphs, p-values were calculated using Fisher’s exact test for all data. The unpaired t-test was used to calculate p-values for comparisons between two groups, while one-way ANOVA with multiple comparisons was used to calculate p-values for analyses involving more than two groups. p-values are NS: not significant, *: p<0.05, **: p<0.01, ***: p<0.001, ****: p<0.0001. GraphPad Prism statistical analysis software and RStudio software were used to generate graphs and perform statistical analyses.

## Supporting Information

This article contains supporting information (45–47).

## Acknowledgements

We thank the Kyoto Stock Center, the Bloomington *Drosophila* Stock Center, the National Institute of Genetics Fly Stocks, and the Vienna *Drosophila* Resource Center for providing fly stocks. We thank Dr. Norbert Perrimon for providing *UAS-BirA*G3-ER*(2), Dr. Ronald P. Kühnlein for providing *FB-Gal4* fly, Dr. Richard W. Carthew for providing *UAS-lacZ-RNAi,* and Dr. C. Goodman for providing *UAS-lacZ*. The proteomics analysis was supported by N. Shinoda.

This work was supported by grants from AMED-Project for Elucidating and Controlling Mechanisms of Aging and Longevity to M.M. under grant no. JP21gm5010001; Japan Society for the Promotion of Science to M.M. under grant numbers 21H04774, 23H04766, 24H00567, and 25H01842; and to S.K. under grant number 21K15100. This work was also supported by the Chugai Foundation for Innovative Drug Discovery Science to S.K. and by the Japan Society for the Promotion of Science and JST SPRING, Grant Number JPMJSP2108 to Y.Y.

We thank the members of the Miura laboratory for their technical assistance and discussions, especially K. Takenaga for preparing the fly food, and N. Shinoda and Y. Nakajima for their helpful discussions and comments.

## Author contributions

Y. Y., S. K., and M. M. designed the experiments and wrote the manuscript. S. K. and M. M. supervised the study. Y.Y. performed the experiments and analyzed the data.

## Declarations

The authors declare no competing or financial interests.

## Notes

### Competing Interest Statement

The authors have declared no competing interest.

### Summary of Updates

Major changes: We have added the following new data as suggested by the reviewers. Added an experiment with a larger sample size due to the high variability in the initial results for Wg morphology (Fig. 1E, F) and Mgat2 knockdown efficiency (Fig. S2A). Confirmed knockdown efficiency of Mgat1 in the FB (Fig. S2B). Investigated the Gnmt-T2A-Gal4 and FBGal4 expression patterns using UAS-GFP (Fig. S2C, D). Presented an overview of the Mgat2-TurboID construct (Fig. 2A) and workflow for secretome analysis (Fig. 2B). Added a list of candidate factors identified by proteomic analysis (Fig. 3D' and Fig. S5D'). Validated Mgat2-TurboID data using various Gal4 drivers other than Gnmt-T2A-Gal4 (Fig. S4B, E, F). Demonstrated that known major secreted factors, such as Lsp1 and Lsp2, were strongly labeled by Mgat2-TurboID (Fig. S5C) to evaluate labeling efficiency of the secreted proteins using Mgat2-TurboID. The fly food table has been moved to the Materials and Methods section to reduce the supplementary tables. Additionally, we have revised the internal descriptions in Table S6 (fly genotype details) for clarity. The updated version is now Table S5. In this revision, we have carefully reviewed the manuscript with a focus on consistent and appropriate terminology. We have revised and refined the wording throughout the text to enhance scientific accuracy and readability.

