## Supplemental Information for "Proximal Labeling of the Golgi Secretome Reveals Fat Body-Derived Humoral Factors in *Drosophila* Disc Regeneration"

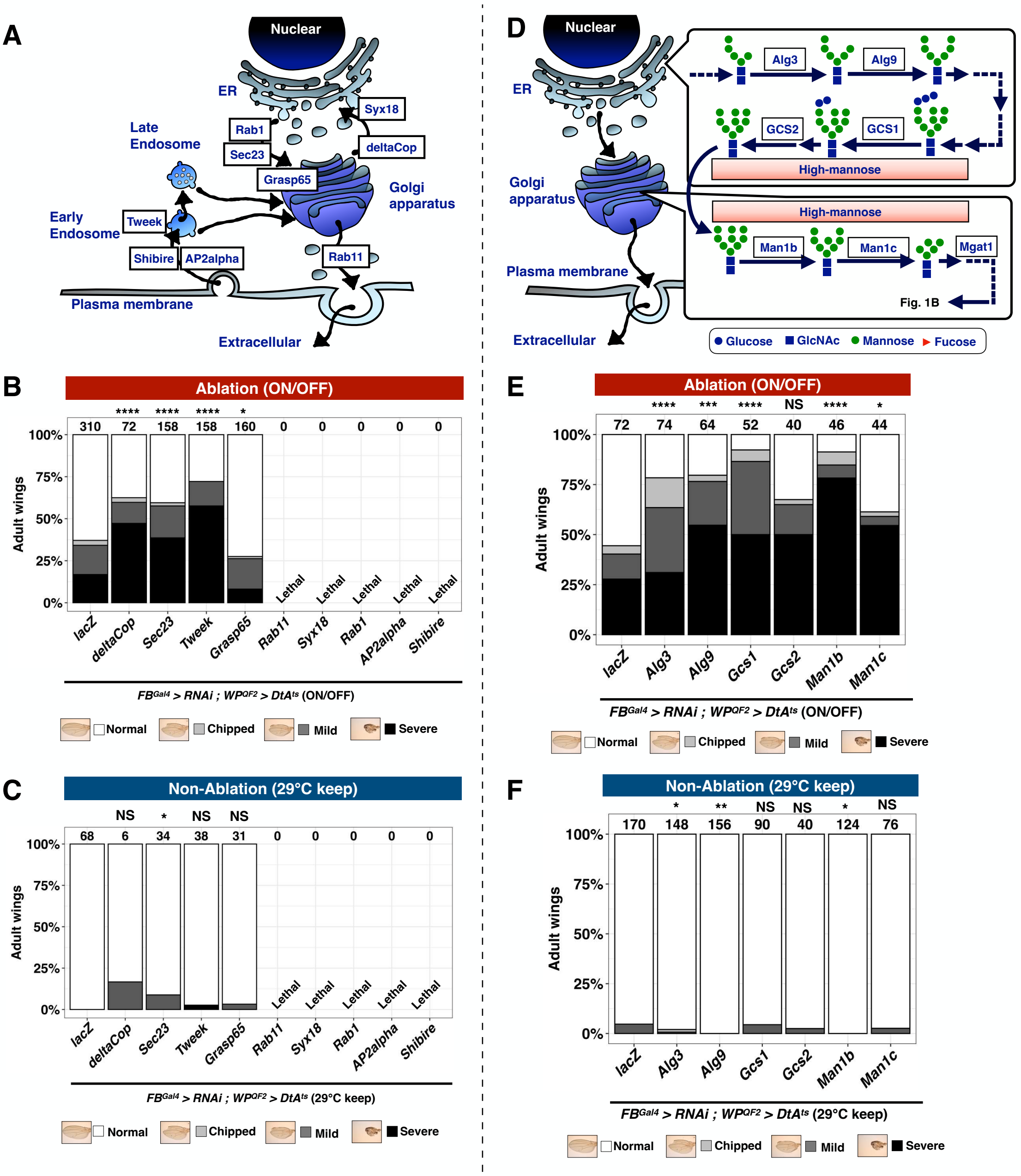

**Figure S1 Requirement of general secretory components in FB for disc regeneration**

(A) Protein trafficking pathway, including secretory pathway and endocytosis.

(B and C) Comparison of adult wing sizes between ablation (B) and non-ablation (C). RNAi knockdown of protein trafficking components hampered disc regeneration. RNAi knockdown was induced in FB by $FB^{Gal4}$. Statistical analysis was conducted using Fisher's exact test to compare control ($WP^{QF2}$>$DtA^{ts}$; $FB^{Gal4}$>$lacZ^{RNAi}$) with treated larvae. NS: not significant, *: p<0.05, ****: p<0.0001. Number of flies is listed above each genotype in the bar graph.

(D) Pathway of N-glycosylation enzymes, including ER. Detailed sequential steps after Mgat1 in the Golgi apparatus were shown in Fig. 1B.

(E and F) Comparison of adult wing sizes between ablation (E) and non-ablation (F). RNAi knockdown of N-glycosylation enzymes hampered disc regeneration. RNAi knockdown was induced in FB by $FB^{Gal4}$. Statistical analysis was conducted using Fisher's exact test to compare control ($WP^{QF2}$>$DtA^{ts}$; $FB^{Gal4}$>$lacZ^{RNAi}$) with treated larvae. NS: not significant, *: p<0.05, **: p<0.01, ***: p<0.001, ****: p<0.0001. Number of flies is listed above each genotype in the bar graph.

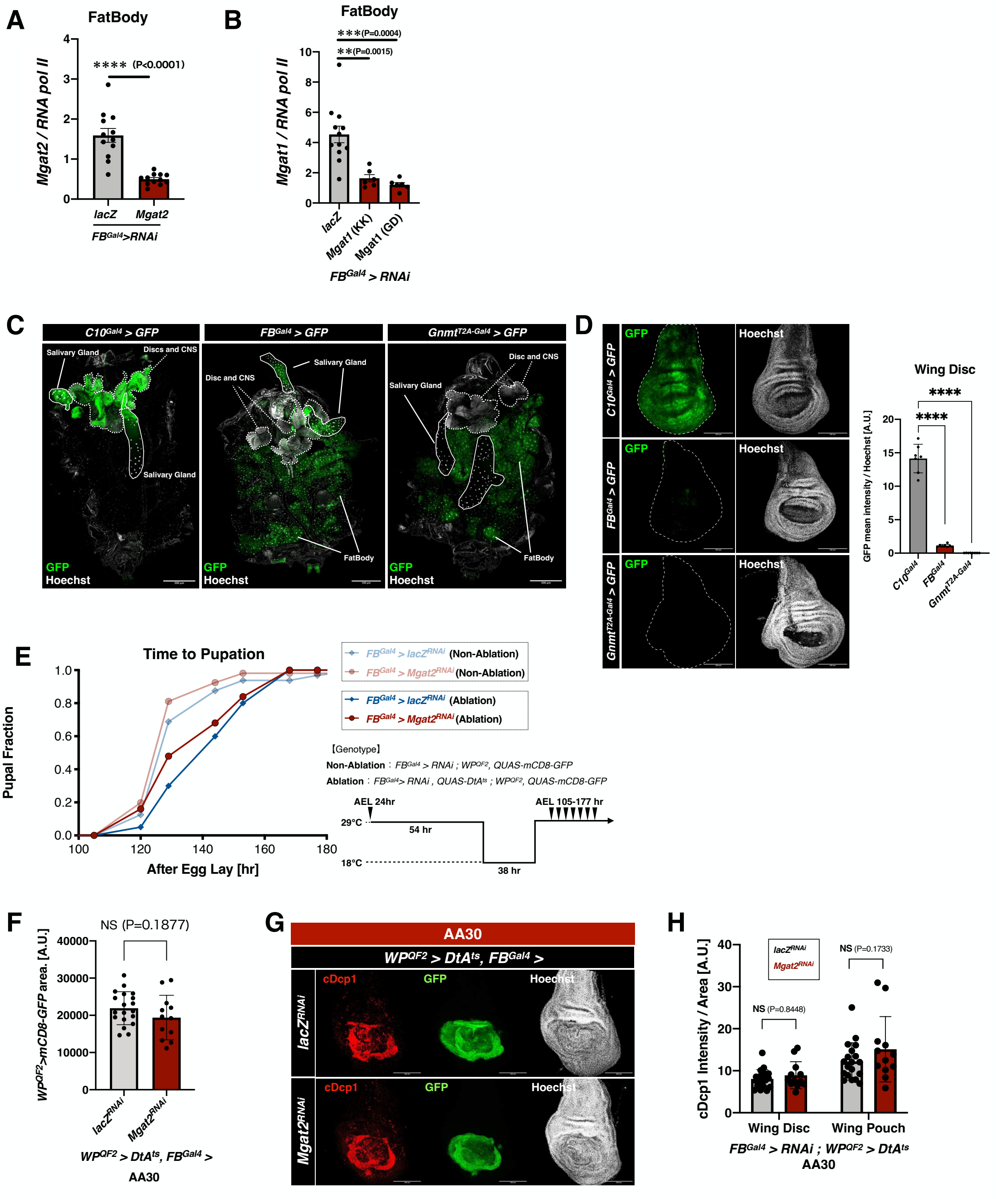

**Figure S2 Effects of Mgat2 knockdown on development and cell death.**

(A) Expression level of *Mgat2* in non-ablated larval fat body. RNAi knockdown of *Mgat2* in FB using $FB^{Gal4}$. mRNA expression was normalized with *RNA pol II* expression. An unpaired t-test was applied. ****: p<0.0001. Number of biological replicates is n=12.

(B) *Mgat1* expression level in the non-ablated larval fat body. RNAi knockdown of *Mgat1* in FB using $FB^{Gal4}$. Each mRNA expression was normalized with *RNA pol II* expression. An unpaired t-test was applied. **: p<0.01, and ***: p<0.001. Number of samples was n=12 in $lacZ^{Gal4}$ and n=6 in $Mgat1^{RNAi}$.

(C, D) Gal4 expression pattern examined by using *UAS-GFP* with FB specific Gal4, $FB^{Gal4}$, and $Gnmt^{T2A\text{-}Gal4}$ (45). $C10^{Gal4}$ as a control of wing disc, salivary gland, and neuron driver. Gal4 expression in whole larvae was shown in (C), and wing disc was shown in (D). GFP was indicated as green, and Hoechst was indicated as white. Number of samples was n=2 in $FB^{Gal4}$ and $Gnmt^{T2A\text{-}Gal4}$, and n=1 in $C10^{Gal4}$ in (C). Number of wing discs is n=8 in $FB^{Gal4}$ and $Gnmt^{T2A\text{-}Gal4}$, and n=7 in $C10^{Gal4}$ in (D).

(E) Pupal fraction for $FB^{Gal4}$ > $Mgat2^{RNAi}$ flies in ablation and non-ablation. n = 32 (NA of $lacZ^{RNAi}$), 106 (NA of $Mgat2^{RNAi}$), 20 (Ablation of $lacZ^{RNAi}$), and 25 (Ablation of $Mgat2^{RNAi}$), respectively, from top to bottom. Temperature treatment was similar between ablation and non-ablation.

(F) $WP^{QF2}$>*mCD8-GFP* area of $FB^{Gal4}$ > $lacZ^{RNAi}$ *and* $Mgat2^{RNAi}$ at AA30 was quantified. Error bars indicate standard error of the mean. An unpaired t-test was applied. NS: not significant. Number of wing discs was n=19 in $lacZ^{RNAi}$ and n=12 in $Mgat2^{RNA}$.

(G) $WP^{QF2}$ > $DtA^{ts}$ induced anti-cleaved Dcp1 (cDcp1) signal in ablated wing pouch region at AA30. White scale bar, 100 μm. Number of wing discs was n=19 in $lacZ^{RNAi}$ and n=12 in $Mgat2^{RNA}$.

(H) cDcp1 intensity in the wing pouch region and whole wing disc region in (G) was quantified. cDcp1-signal amount was unchanged between $FB^{Gal4}$ > $lacZ^{RNAi}$ and $FB^{Gal4}$ > $Mgat2^{RNAi}$. Error bars indicate standard error of the mean. An unpaired t-test was applied. NS: not significant. Number of wing discs was n=19 in $lacZ^{RNAi}$ and n=12 in $Mgat2^{RNA}$.

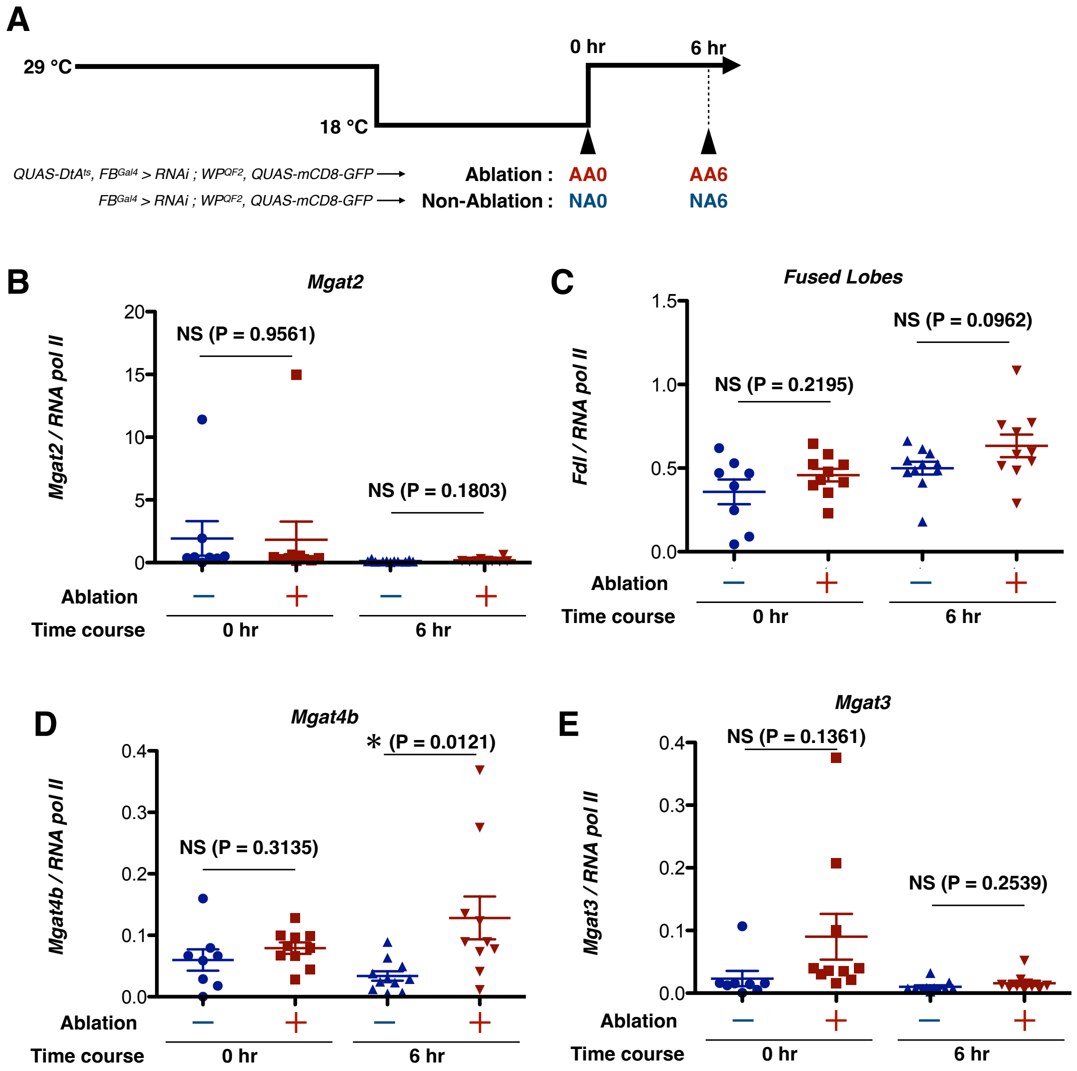


**Figure S3 Expression of glycosyltransferases in FB during disc regeneration.**

(A) Scheme of the sampling time course. Sampling was performed at two time points: 0 and 6 h after ablation (AA). Ablation or Non-Ablation (NA) conditions were applied to DtA$^{ts}$-expressing or non-DtA$^{ts}$-expressing larvae, respectively, and they were maintained under the same temperature shift.

(B-E) *Mgat2*, *Fused Lobes*, *Mgat4,* and *Mgat3* expression level in the disc-ablated and non-ablated larval fat body. Each mRNA expression was normalized with *RNA pol II* expression. One-way ANOVA and Tukey's multiple comparison test were applied. NS: not significant, *: $p<0.05$. Number of samples was n=8 in NA0, n=10 in AA0, n=11 in NA6, and n=10 in AA6.

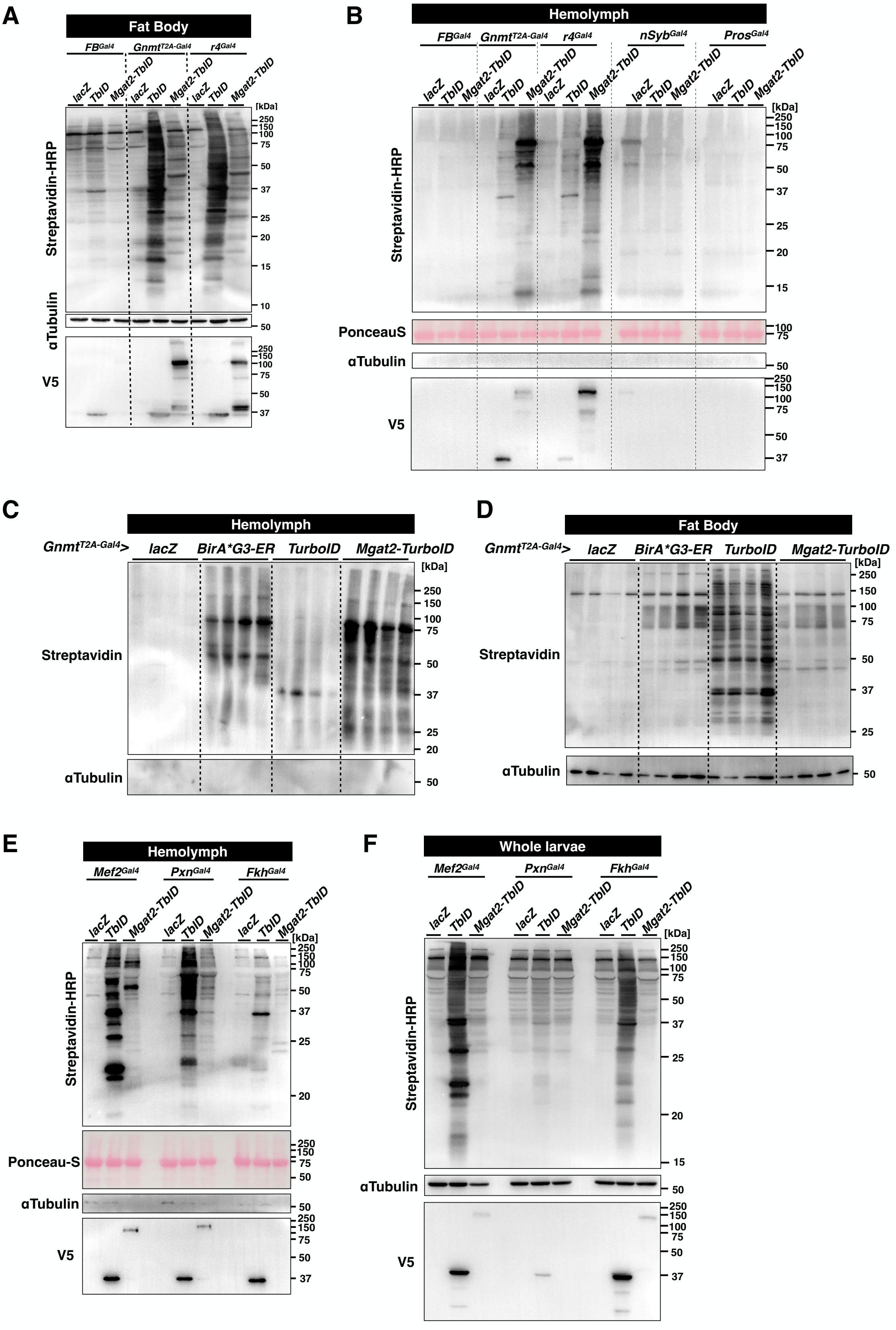


**Figure. S4 Validation of secretory protein labeling with Mgat2-TurboID.**

(A and B) Western blotting for biotinylated proteins (detected by Streptavidin-HRP) in FB (A) and hemolymph (B). *LacZ*, *TurboID,* and *Mgat2-TurboID* were expressed in FB with $FB^{Gal4}$, $Gnmt^{T2A\text{-}Gal4}$, $r4^{Gal4}$, in neurons with $nSyb^{Gal4}$, and neural cells and gut enteroendocrine cells with $Pros^{Gal4}$. Samples were collected in Non-Ablation (NA) condition. αTubulin and Ponceau-S were used as an internal control, and anti-V5 antibody was used to detect expression and secretion of the constructs. 5–10 larvae were used for each lane.

(C and D) Western blotting for biotinylated proteins (detected by Streptavidin-HRP) in hemolymph (C) and FB (D). *LacZ*, *BirA*G3-ER*, *TurboID*, and *Mgat2-TurboID* were expressed in FB with $Gnmt^{T2A\text{-}Gal4}$. Samples were collected in Non-Ablation (NA) condition. αTubulin was used as an internal control. 10–20 larvae were used for each lane.

(E and F) Western blotting for biotinylated proteins (detected by Streptavidin-HRP) in hemolymph (E) and FB (F). *LacZ*, *TurboID*, and *Mgat2-TurboID* were expressed in muscle with $Mef2^{Gal4}$, in hemocyte with $Pxn^{Gal4}$, and salivary gland with $Fkh^{Gal4}$. Samples were collected in Non-Ablation (NA) condition. αTubulin and Ponceau-S were used as an internal control, and anti-V5 antibody was used to detect expression and secretion of the constructs; 5–10 larvae were used for each lane.

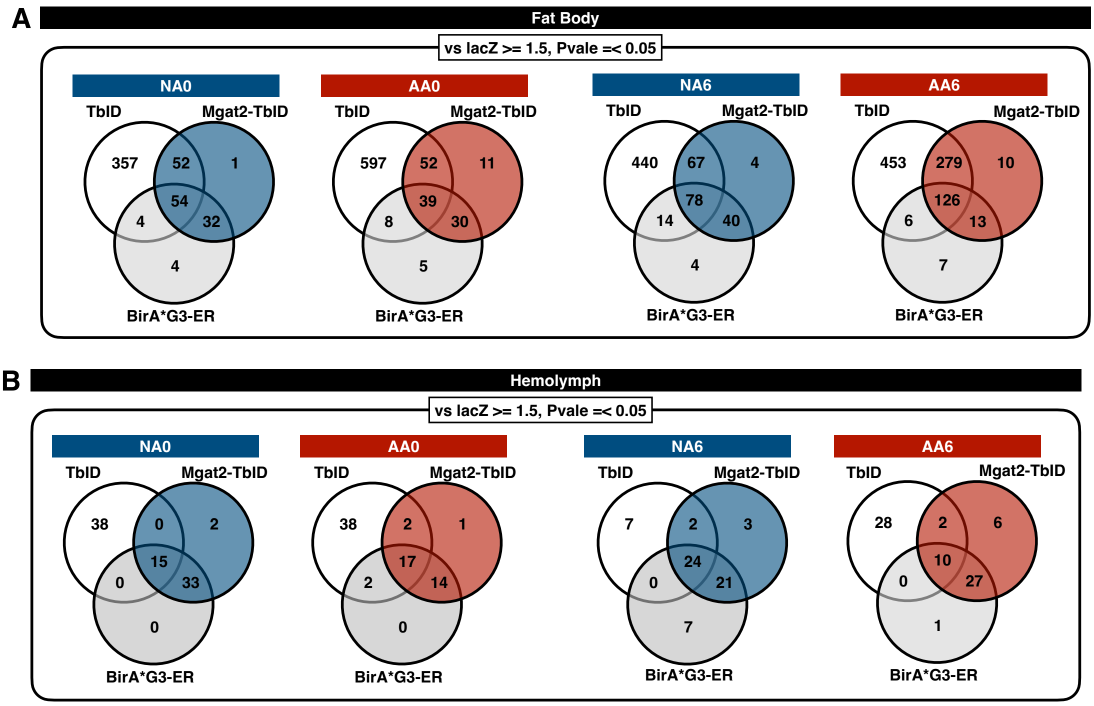


C

| Hemolymph sample | Abundance of Mgat2-TbID / lacZ | | | | Abundance of V5-TbID / lacZ | | | |
|---|---|---|---|---|---|---|---|---|
| Protein Name | NA0 | AA0 | NA6 | AA6 | NA0 | AA0 | NA6 | AA6 |
| Larval serum protein 1α | 100 | 100 | 100 | 100 | 2.707 | 4.898 | 9.786 | 3.278 |
| Larval serum protein 1γ | 100 | 100 | 94.268 | 73.248 | 1.432 | 2.407 | 2.563 | 1.494 |
| Larval serum protein 1β | 100 | 100 | 100 | 100 | 2.970 | 2.209 | 3.105 | 2.267 |
| Larval serum protein 2 | 100 | 100 | 100 | 100 | 2.285 | 3.319 | 10.791 | 1.961 |

E

| GO term (BP Direct) | | Count | PValue |
|---|---|---|---|
| response to fungus | Immune | 4 | 1.88E-07 |
| iron ion transport | Iron Homeostasis | 3 | 3.96E-05 |
| detoxification of iron ion | | 2 | 1.79E-03 |
| iron ion import across plasma membrane | | 2 | 1.79E-03 |
| intracellular sequestering of iron ion | | 2 | 2.69E-03 |
| cellular iron ion homeostasis | | 2 | 0.0125 |

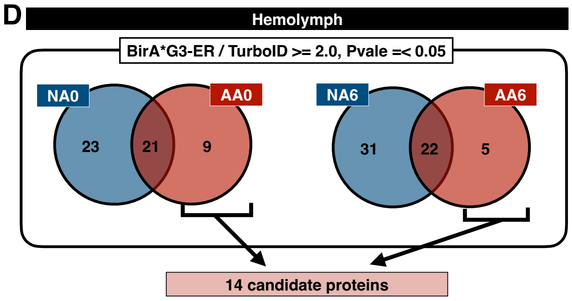


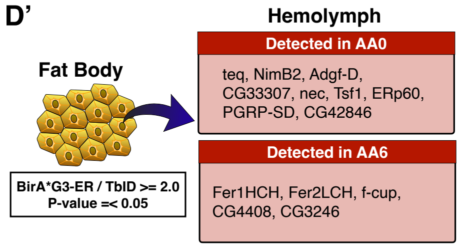


**Figure. S5 Proximal proteins labeled by Mgat2-TurboID closely resemble those labeled BirA*G3-ER.**

(A and B) Analysis of biotinylated proteins in hemolymph (A) and FB (B) at two time points. Proteins detected with each genotype at ≧1.5 times compared to *lacZ* and a p-value of ≤0.05 were extracted, and their numbers were listed.

(C) Labeled abundance of major larval secretory proteins (Lsp1α, Lsp1β, Lsp1γ, and Lsp2) in the hemolymph sample. Abundance ratio of Mgat2-TurboID vs lacZ and TurboID vs lacZ was used. In the analysis by Proteome Discoverer 2.2, ratio abundance data displays a maximum value of 100, so values greater than or equal to 100 are displayed as 100.

(D and D') Analysis of biotinylated proteins in the hemolymph at two time points. Under each condition, proteins detected with *BirA*G3-ER* at more than twice the amount when compared to *V5-TurboID*, and with a p-value of 0.05 or less, were extracted. Among these proteins, 14 were specific to Ablation (D'). 15–25 larvae were used for a single sample, and biological replicate was n=3.

(E) GO analysis of 14 proteins in (D) with DAVID.
